# Aurora B/C-dependent phosphorylation promotes Rec8 cleavage in mammalian oocytes

**DOI:** 10.1101/2021.10.22.465153

**Authors:** Elvira Nikalayevich, Safia El Jailani, Damien Cladière, Yulia Gryaznova, Célia Fosse, Sandra A. Touati, Eulalie Buffin, Katja Wassmann

## Abstract

To generate haploid gametes, cohesin is removed in a step-wise manner from chromosome arms in meiosis I and the centromere region in meiosis II, to segregate chromosomes and sister chromatids, respectively. Meiotic cohesin removal requires cleavage of the meiosis-specific kleisin subunit Rec8 by the protease Separase[1, 2]. In yeast, Rec8 is kept in a non-phosphorylated state by the action of PP2A-B56, which is localised to the centromere region, thereby preventing cohesin removal from this region in meiosis I[3-5]. However, it is unknown whether Rec8 has to be equally phosphorylated for cleavage, and whether centromeric cohesin protection is indeed brought about by dephosphorylation of Rec8 preventing cleavage, in mammalian meiosis. The identity of one or several potential Rec8-specific kinase(s) is also unknown. This is due to technical challenges, as Rec8 is poorly conserved preventing a direct translation of the knowledge gained from model systems such as yeast and *C. elegans* to mammals, and additionally, there is no turn-over of Rec8 after cohesion establishment, preventing phosphomutant analysis of functional Rec8. To address how Rec8 cleavage is brought about in mammals, we adapted a biosensor for Separase to study Rec8 cleavage in single mouse oocytes by live imaging, and identified phosphorylation sites promoting cleavage. We found that Rec8 cleavage by Separase depends on Aurora B/C kinase activity, and identified a residue promoting cleavage and being phosphorylated in an Aurora B/C kinase-dependent manner. Accordingly, inhibition of Aurora B/C kinase during meiotic maturation impairs endogenous Rec8 phosphorylation and chromosome segregation.

## Results and Discussion

Yeast Rec8 is phosphorylated by CK1δ/*ε* (Hrr25 in *S. cerevisiae*, Hhp2 in *S. pombe*) and DDK (Cdc7-Dbf4 in *S. cerevisiae*, possibly Hsk1/Dfp1 in *S. pombe*) for cleavage by Separase[6-9], whereas in *C. elegans* oocytes, Rec8 is phosphorylated by AIR-2 (Aurora B), which is recruited to the short chromosome arms in a Haspin kinase-dependent manner to bring about Rec8 cleavage[10]. We hypothesized that also in vertebrate meiosis one of these kinases may be required for Rec8 cleavage.

Aurora kinases are primarily regulated through their localization and interaction with their molecular partners, and not their individual substrate specificities. In mammalian oocytes three Aurora kinases are expressed, namely Aurora A, B and the oocyte-specific Aurora C[11, 12]. To ask whether chromosome segregation defects indicative of failure to cleave Rec8 are observed upon loss of both Aurora B and C, oocytes entering meiosis I were treated with two inhibitors targeting both Aurora B and C, AZD1152 and ZM447439 [13-17]. Efficient Aurora B/C inhibition was verified by staining metaphase I chromosome spreads with a phospho-specific antibody against Histone H3pS10, a prominent substrate of both kinases also in oocytes[15, 16, 18] (**Figure 1a**). Because of Aurora B/C’s role in cytokinesis preventing PB (Polar body) extrusion [16, 17, 19], we verified whether these oocytes nevertheless underwent metaphase-to-anaphase transition, a condition to analyse chromosome segregation. Exogenously expressed Securin-YFP degradation was assessed by live imaging and as previously published, occurred in an accelerated manner in oocytes without Aurora B/C activity due to the absence of error correction signaling to the Spindle Assembly Checkpoint (SAC) (**Figure 1b**)[17, 19]. Securin-YFP degradation shows that inhibitor-treated oocytes are not cell-cycle arrested and proceed through meiosis I. To determine whether bivalent chromosomes segregate in absence of Aurora B/C kinase activity, chromosome spreads were prepared from metaphase II oocytes treated with Aurora B/C inhibitors throughout meiotic maturation. Whereas control oocytes harbored only paired sister chromatids (dyads), the majority of oocytes treated with Aurora B/C inhibitors contained several bivalents (paired chromosomes), indicating failures in meiosis I chromosome segregation (**Figure 1c-g**). This phenotype is not due to absence of error correction resulting in chromosomes being attached to only one pole, because -as we have shown previously-bivalents attached to a monopolar spindle still separate homologous chromosomes[20]. Hence, Aurora B/C kinase may be involved in efficient arm cohesin removal in meiosis I.

**Figure 1.**
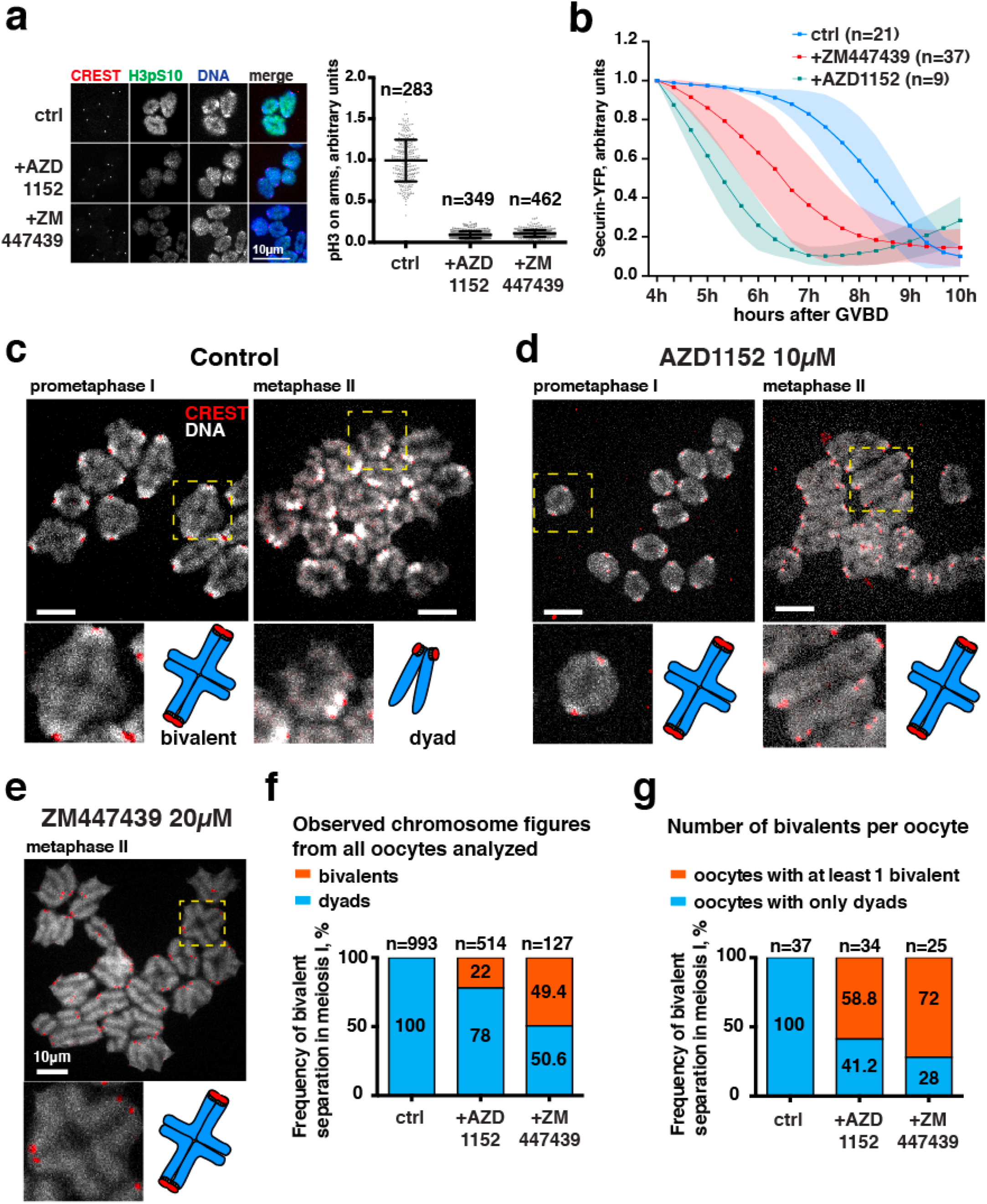
Aurora B/C activity is required for homologous chromosome separation in meiosis I. **a)** Chromosome spreads from mouse oocytes in prometaphase I (GVBD+4h) stained for DNA (with Hoechst, blue), centromeres (with CREST antibody, red), and H3 phosphorylated on Ser10 (H3pS10, green). Scale bar (white) is 10 *μ*m. The graph on the right shows the corresponding quantification of H3pS10 fluorescence intensity detected on chromosome arms. Each dot is a measurement from one chromosome arm corrected for background. Bars and whiskers are means and SD. **b)** Accelerated Securin-YFP degradation in oocytes treated with AZD1152 or ZM447439. Securin-YFP fluorescence intensity in the cytoplasm was measured in time lapse images of mouse oocytes undergoing maturation. n is the number of oocytes. Hours after Germinal Vesicle Breakdown (GVBD) are indicated. Time points were taken every 20 minutes. Each dot and whiskers (shaded areas) are means and SD. **c)** Chromosome spreads from mouse oocytes in prometaphase I (GVBD+4h) and metaphase II (GVBD+18h), stained for DNA (with Hoechst, grey) and centromeres (with CREST antibody, red). Lower panels are enlarged areas marked with yellow dash line squares and a schematic of a bivalent (left) and a sister chromatid pair (right). On the schematic the chromosome arms are in blue and centromeres are in red. Scale bar (white) is 10 *μ*m. **d)** Chromosome spreads from mouse oocytes treated with 10 *μ*M Aurora B/C inhibitor AZD1152 from GVBD onwards, in prometaphase I (GVBD+4h) and metaphase II (GVBD+18h), stained for DNA (with Hoechst, grey) and centromeres (with CREST antibody, red). Lower panels are enlarged areas marked with yellow dash line squares and a schematic of bivalents. On the schematic the chromosome arms are in blue and centromeres are in red. Scale bar (white) is 10 *μ*m. **e)** Chromosome spreads from mouse oocytes treated with 20 *μ*M Aurora B/C inhibitor ZM447439 from GVBD onwards, in metaphase II (GVBD+18h), stained for DNA (with Hoechst, white) and centromeres (with CREST antibody, red). Lower panels are an enlarged area as marked with a yellow dash line square and a schematic of a bivalent. On the schematic the chromosome arms are in blue and centromeres are in red. Scale bar (white) is 10 *μ*m. **f)** Frequency of bivalent separation in meiosis I, quantified from chromosome spreads of metaphase II oocytes (GVBD+18h). Chromosomes were observed in control oocytes and oocytes treated with Aurora B/C inhibitors (10 *μ*m AZD1152 or 20 *μ*M ZM447439) and classified as “bivalents” (homologous chromosomes together in form of a bivalent), or “dyads” (homologous chromosomes separated into sister chromatid pairs). The number for dyads is divided by 2 to compare to the number of “not separated bivalents”. n is the number of analyzed bivalents. *** p<0.001, calculated with Fisher’s exact test. **g)** Same data as f), but numbers are per oocyte. Only groups of chromosomes for one oocyte in the same field were quantified. n is the number of oocytes. On each graph all frequencies are displayed as numbers on the corresponding bars. *** p<0.001, calculated with Fisher’s exact test. Related to Figure S1.

Loss of Aurora B/C can be complemented in part by Aurora A[21], and complete loss of Aurora A interferes with proper spindle formation[22, 23], leading to a cell cycle arrest in metaphase I[23]. To address whether Aurora A is also required for chromosome segregation in meiosis I, we treated oocytes with the Aurora A inhibitor MLN8237[24] from entry into meiosis I onwards. Such as Aurora A knock-out oocytes, MLN8237-treated oocytes arrested in metaphase I, indicating that inhibition of Aurora A was efficient[22, 23] (**Figure S1a**). We added the Mps1 inhibitor Reversine to override the SAC arrest[25] which is at least partially responsible for the metaphase I arrest induced by Aurora A inhibition[23], and performed chromosome spreads 14 hours later. **Figure S1a** shows that inhibition of Aurora A and simultaneously overriding the SAC arrest with Reversine rescued PB extrusion in 40 % of MLN8237 treated oocytes, similar to previously published data[23]. Importantly, MLN8237 treated oocytes that extruded a PB also segregated chromosomes (**Figure S1b**). Hence, we conclude that Aurora A is not required for cohesin removal in oocyte meiosis I.

It was attractive to speculate that Aurora B/C directly phosphorylate Rec8 to convert it into a Separase substrate, but it was just as likely that Aurora B/C activity were required for Separase activation itself, independently of APC/C (Anaphase Promoting Complex/Cyclosome) activation. Separase activity can be analysed in live cells using a biosensor containing the mitotic kleisin subunit Rad21 sandwiched between two fluorescent tags and fused to H2B, to localize the sensor to chromosomes for quantification [26, 27]. Of note, in oocytes expression of this and similar constructs containing kleisin subunits does not produce cohesive cohesin complexes, as those are put in place long before entry into the first meiotic division, before birth of the female, and there is no turnover during the time of expression upon mRNA injection at GV (germinal vesicle) stage[28]. Loss of co-localization of the two fluorophores on chromosomes indicates Separase cleavage of the sensor (**Figure 2a**). Notably, sensor cleavage was not perturbed by addition of any of two Aurora B/C kinase inhibitors from entry into meiosis I onwards (**Figure 2b**), demonstrating that Separase is activated at the right time and with comparable efficiency in the presence or absence of Aurora B/C kinase activity. Aurora B/C kinases are therefore not required for efficient Separase activation in meiosis I.

**Figure 2.**
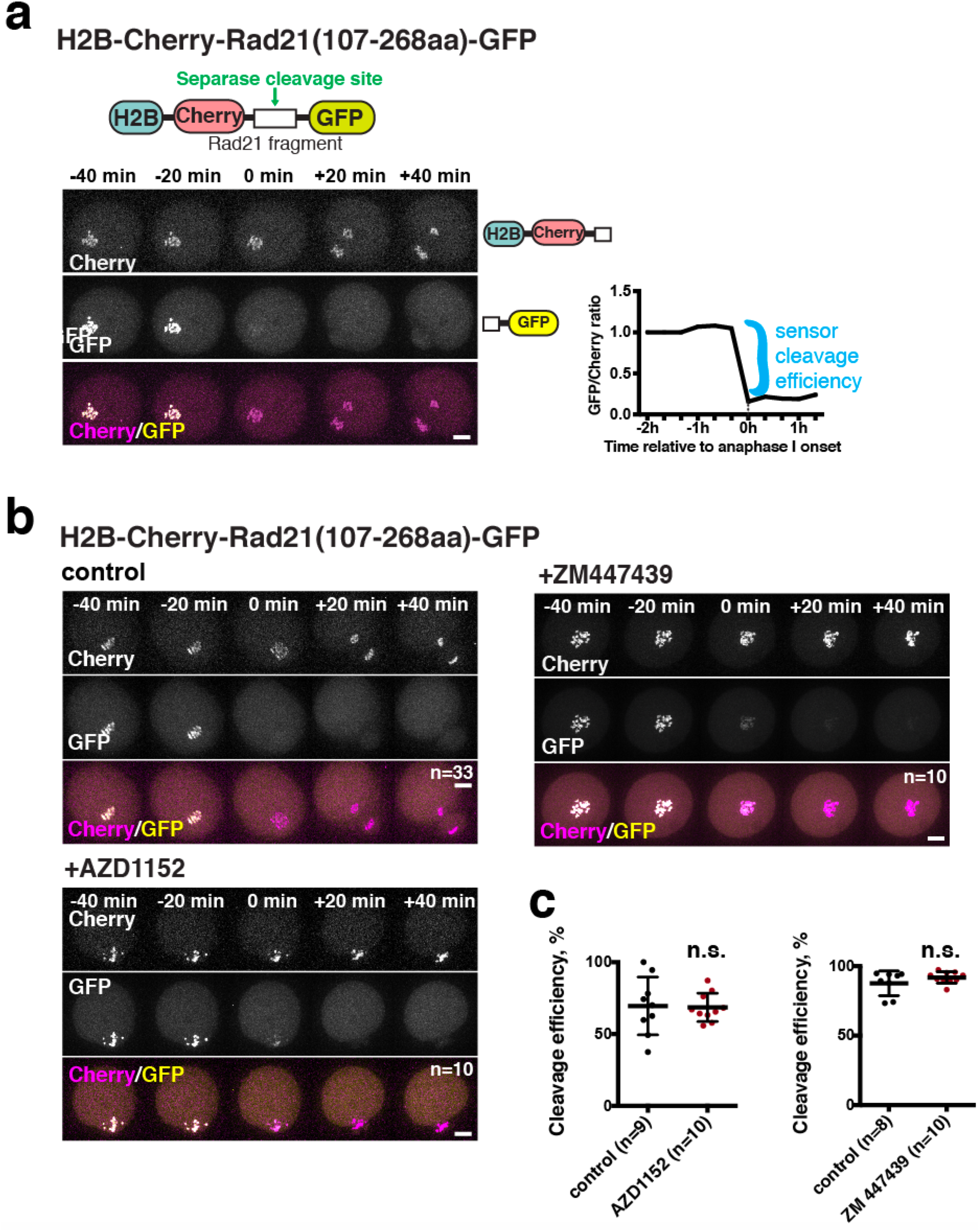
Aurora B/C activity does not inhibit Separase-dependent cleavage of Rad21 sensor. **a)** Schematic of sensor constructs (top), example of time lapse microscopy acquisition of a mouse oocyte expressing the Rad21 sensor, +/- 40 minutes relative to anaphase I onset (middle) and an example of the quantification of GFP/Cherry ratio (on the right). Anaphase I is labeled as 0 minutes timepoint. Scale bar (white) is 20 *μ*m. Cleavage of the sensor is visible by the disappearance of the GFP signal from the chromosomes, while the Cherry signal remains. Quantification of the GFP/Cherry ratio measures mean signal from the area occupied by the chromosomes corrected for the background signal from the cytoplasm. Sensor cleavage efficiency is quantified as per cent decrease of the GFP/Cherry ratio when comparing the last timepoint before anaphase I onset and the lowest point during anaphase I. **b)** Cleavage of Rad21 sensor in control oocytes and oocytes treated with the indicated inhibitors of Aurora B/C. Shown are examples of time lapse microscopy images of mouse oocytes expressing the Rad21 sensor, +/- 40 minutes relative to onset of anaphase I. Scale bar (white) is 20 *μ*m. **c)** Quantification of cleavage efficiency of Rad21 sensor from experiment in b). n is the number of oocytes analysed. Bars and whiskers are mean and SD. n.s. – not significant, according to Mann-Whitney test.

To ask whether Aurora B/C targets Rec8 for cleavage by Separase, we decided to adapt the Rad21 sensor assay to Rec8. Full-length Rec8 was cloned into the sensor construct instead of the Rad21 fragment and *in vivo* cleavage by Separase was analysed by quantifying loss of co-localization of the two fluorophores in oocytes. A recent study had shown that a Rec8 sensor spanning aa 297-506 is not cleaved *in vivo*, with the caveat that cleavage assays were performed in mitotic cells[29]. In our hands, full-length Rec8 sensor was cleaved *in vivo* in meiosis I oocytes, albeit with low efficiency (**Figure 3a)**. Importantly though, this cleavage was abolished when 39 out of 59 Serines and Threonines were mutated to Alanine (**Figure 3a, Table S1)**, indicating that phosphorylation of at least some of those sites is required for Rec8 cleavage.

**Figure 3.**
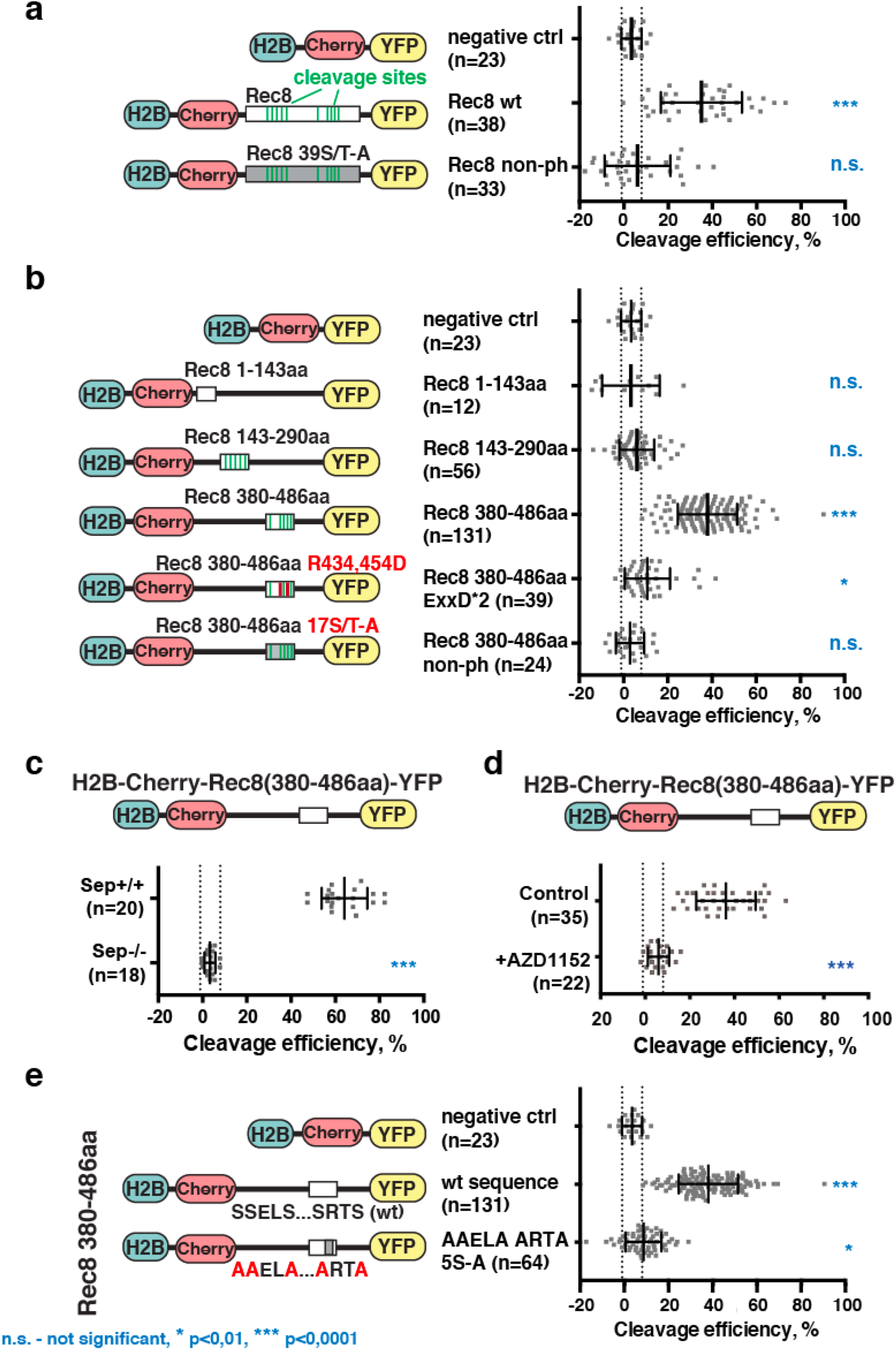
Cleavage efficiencies of Rec8 sensor constructs in oocyte meiosis I. The sensor constructs that were analysed are indicated on the left with corresponding schemes, and the corresponding *in vivo* cleavage efficiencies when expressed in oocytes, are on the right. Negative control (ctrl) indicates a sensor construct without Rec8 sequence and no cleavage site. In the schemes, potential Separase cleavage sites are indicated in green, single amino acid substitutions in red, multiple amino acid substitutions indicated in grey, and the relative position of the Rec8 fragment being tested is shown by its position on the black line (not in scale). On each graph n is the number of oocytes, bar and whiskers are means and SD. Vertical dotted lines mark standard deviation of the negative control sensor. n.s. – not significant, * p<0.01, *** p<0.0001 according to t test performed for each condition compared to negative control. **a)** Cleavage of sensor construct containing full-length Rec8, and Rec8 harboring 39 Serine or Threonine mutated to Alanine. **b)** Cleavage efficiencies of three Rec8 fragments (1-143aa, 143-290aa, 380-486aa), and Rec8 380-486aa harboring mutations in two Separase cleavage sites, or 17 Serine or Threonine mutated to Alanine. **c)** Cleavage efficiencies of Rec8 380-486aa sensor construct in Separase control (Sep^+/+^) and Separase knock-out (Sep^-/-^) oocytes. **d)** Cleavage efficiencies of Rec8 380-486aa sensor construct with or without being exposed to Aurora B/C inhibitor AZD1152 from GVBD onwards. **e)** Cleavage efficiencies of Rec8 380-486aa harboring Alanine substitutions in five potential phosphorylation sites (S467, S470, S478, S479, and S482), S482 corresponding to an Aurora B consensus site. Related to supplementary Figure S2. See also supplementary Table 1.

The low cleavage efficiency of the full-length sensor prompted us to test truncated versions of Rec8 in our sensor assay, to reduce potential interference of other cohesin binding proteins when examining phosphorylation site mutations. Not surprisingly, a Rec8 fragment spanning the first 143 aa and not containing any potential Separase cleavage sites was not cleaved at all in oocytes. A Rec8 fragment spanning 143-290 aa was not cleaved either, even though it contained five potential minimal Separase cleavage sites, one of which allowed Rec8 cleavage *in vitro* in a previous study[30]. The two most likely Separase cleavage sites (aa 429 and 449)[30] are found in Rec8 fragment 380-486aa, and indeed, this fragment was efficiently cleaved in our sensor assay (**Figure 3b, Figure S2a)**. Cleavage did not occur upon mutation of both prominent Separase cleavage sites (Rec8 380-486aa ExxD*2). (**Figure 3b, Table S1)**. Accordingly, cleavage was Separase dependent, because it did not take place in oocytes devoid of Separase due to a conditional knock-out [31] (**Figure 3c, Figure S2b)**. Therefore, in oocytes a biosensor containing a Rec8 fragment can be used to dissect the requirements for Separase cleavage, unlike expression of a truncated sensor in mitotic cells, which needs to be fused with the protein Meikin, to be cleaved [29].

Cleavage of the C-terminal Rec8 380-486aa fragment was abolished when 17 Serines and Threonines **(Table S1)** were mutated to Alanine, indicating that phosphorylation of at least some of these residues is necessary for efficient Separase cleavage (**Figure 3b)**. Furthermore, sensor cleavage did not occur in the presence of Aurora B/C inhibitor (**Figure 3d)**. Mutating only 5 Serines to Alanines around a conserved Aurora B/C, CK1 and Plk1 consensus site was enough to abrogate cleavage of the sensor (Rec8 380-486aa 5S-A) (**Figure 3e)**. Altogether, our data strongly indicate that Aurora B/C dependent phosphorylation of Rec8 -directly or indirectly-promotes Separase-dependent cleavage of Rec8.

Our results demonstrate that cleavage of exogenously expressed Rec8 fragments requires certain phosphorylatable residues and does not take place without Separase and Aurora B/C kinase activity. Our data, however, do not yet show that endogenous Rec8 is indeed phosphorylated in oocyte meiosis I in an Aurora B/C-dependent manner. In meiosis I, cohesin on chromosome arms is removed during the first division to allow separation of chromosomes, whereas cohesin in the centromere region is protected to maintain sister chromatids together until meiosis II[1, 2]. Therefore, we expected endogenous Rec8 to be phosphorylated on arms in meiosis I, and in the centromere region in meiosis II. But before addressing this issue with phospho-specific antibodies, we wanted to determine the amount of endogenous centromeric Rec8 present in meiosis I and maintained until meiosis II, to check whether we could actually expect to distinguish distinctly phosphorylated fractions of endogenous Rec8 around the centromere, by immunostaining. We have discovered recently that an event we termed kinetochore individualization takes place in anaphase I and may require cleavage of centromeric Rec8[20]. Indeed, additionally to arm- and pericentromere-localized Rec8, a third fraction at the centromere was recently shown to be removed at exit from meiosis I[32], and may therefore also be modified by phosphorylation already in metaphase I. To address how much Rec8 is effectively protected in the centromere region, we stained endogenous Rec8 with a polyclonal antibody on chromosome spreads, obtained from meiosis I and meiosis II metaphase oocytes. Oocytes were fixed in the same chambers, to best compare the amounts of Rec8 present in the centromere region. Kinetochores and underlying centromeres were stained with CREST serum, and the Rec8 signal overlapping with CREST and in-between the two CREST signals was quantified and normalized. We found that less than 20% of Rec8 present between and overlapping with the two centromere signals is preserved from meiosis I to meiosis II **(Figure 4a)**. In other words, 80 % of Rec8 is removed from this region already in meiosis I and may thus be phosphorylated, similar to Rec8 localized to arms. This result indicates that phospho-specific antibodies directed against the sites we identified can be expected to stain Rec8 on arms as well as in the centromere region in meiosis I.

**Figure 4.**
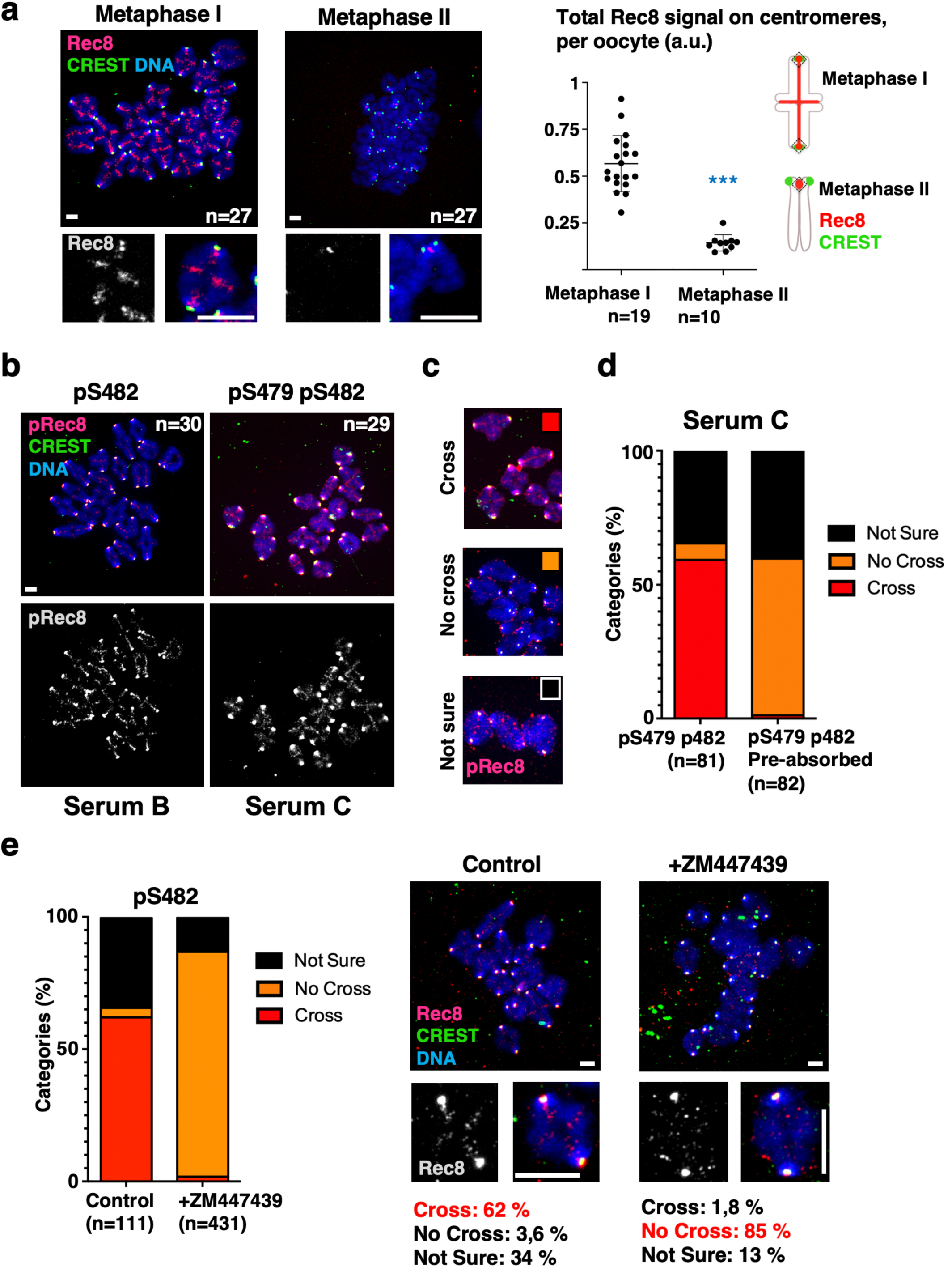
Phospho-specific antibodies recognize Rec8 in an Aurora B/C kinase dependent manner. **a)** Oocytes in metaphase I and metaphase II were fixed for chromosome spreads right next to each other, in the same wells, and stained with a polyclonal Rec8 antibody (magenta), anti-CREST serum (Centromeres, green), and Hoechst (DNA, in blue). A representative spread and a zoom on one bivalent are shown for each condition on the left. n indicates the number of oocytes analysed, scale bars are 10 *μ*m. On the right, quantifications of Rec8 signal overlapping with CREST, or inbetween the two CREST signals, as indicated in the scheme. *** p<0.0001, by Mann-WhitneyTest. **b)** Chromosome spreads of oocytes in prometaphase I were stained with the indicated sera to detect endogenous Rec8 phosphorylated on S482, or both S479 and S482 (magenta), together with anti-CREST serum (Centromeres, green), and Hoechst (DNA, in blue). A representative spread is shown for each condition. n indicates the number of oocytes analysed, scale bars are 10 *μ*m. **c)** Spreads stained with phospho-specific anti-Rec8 antibodies were classified according to the examples of stainings shown in b). **d)** Serum C was pre-absorbed with peptide pS479 pS482, and used to stain chromosome spreads as described in b). The results of quantifications according to the classification in c) are shown. **e)** Chromosome spreads of control oocytes in prometaphase I and oocytes treated with ZM447439 from GVBD onwards, were stained with serum B to detect endogenous Rec8 phosphorylated on S482 (magenta), together with anti-CREST serum (Centromeres, green), and Hoechst (DNA, in blue). The graph shows quantifications of the indicated number of chromosomes, according to classification in c). A representative spread with a zoom on one chromosome is shown for each condition. n indicates the number of oocytes analysed, scale bars are 10 *μ*m. Related to supplementary Figure 3.

We produced phospho-specific antibodies in rabbits against the sites identified with the Rec8 Separase sensor. Each rabbit was primed for one of the two sites and with 3 peptides for a given site (“SELS”, containing S467 and S470; and “SRTS”, containing S479 and S482, see **Figure S3a** for scheme and peptide sequences). S467 when phosphorylated may correspond to a priming site for CK1-dependent phosphorylation of S470, but S470 also corresponds to a Plk1 consensus site[33]. S482 constitutes a potential Aurora B/C as well as CK1 phosphorylation site, whereas pS479 may serve as a priming site and together with pS482 may contribute to creating a powerful phospho-epitope recognized by phospho-specific antibodies.

Aliquots of the sera from the rabbits with the highest titer for each site were purified against each of the three peptides separately and used for immunostaining of chromosome spreads in meiosis I. If the phospho-specific antibodies were specific for the phosphorylated fraction of Rec8 they should recognize the truncated sensor construct in its wild type form (assuming it becomes phosphorylated) but not the corresponding Rec8 380-486aa Serine to Alanine mutants, when expressed in oocytes. This was indeed the case for the antibodies purified against phosphorylated S479 and S482, the potential Aurora B consensus site, alone (for pS482) or in combination, as Sera B and C recognized the Rec8 380-486aa wt but not the Rec8 380-486aa S479, 482A sensor **(Figure S3b,c)**. On the other hand, from two rabbits none of the purified sera against pS467 and pS470, the Plk1 consensus site, again either alone or in combination, recognized the Rec8 380-486aa wt sensor, suggesting that either these Serines are not phosphorylated, or phosphorylation does not create a powerful antigen **(Figure S3b)**. We therefore focused on the SRTS Aurora B consensus motif in the remaining experiments, using exclusively sera B and C.

Antibody sera against pS482 and pS479 pS482 (B, C) detected endogenous arm-localized cohesin in prometaphase I, recognizable by the typical cross-shaped pattern that is usually observed when staining for Rec8 **(Figure 4b)**, and indicating that endogenous Rec8 is phosphorylated on S482 and potentially also on pS479. To assess presence of phosphorylated Rec8 on chromosome arms, we divided our chromosome spreads into three categories, as the intensity of the signal was highly variable, depending on the quality of the individual spreads, the degree of chromosome condensation and could therefore not be quantified **(Figure 4c)**. As negative controls we used either the pre-serum, or the serum pre-absorbed with the SRTS peptide pS479 pS482 **(Figure 4d)**. A prominent signal was also detected in the centromere region with B and C sera **(Figure 4b)**. Centromeres have the potential to generate an unspecific background signal, with multiple Aurora kinase substrates being present there[3] that may cross-react with the serum we generated. To address this issue, we compared chromosome spreads from embryos at the 8-cell stage undergoing mitosis, with meiosis II chromosome spreads. Because Rec8 is meiosis-specific and substituted by Scc1 from the first mitosis onwards[34], this allowed us to distinguish the signal specific for Rec8 on paired sister chromatids in meiosis II from a background signal present at centromeres of paired sister chromatids in mitosis, on samples prepared exactly the same manner. Indeed, an unspecific signal colocalizing with CREST was detected on mitotic and meiotic spreads with serum C purified against both phosphorylation sites, whereas in addition to this unspecific signal, another signal in-between sister centromere dots exactly where centromeric cohesin is localized **(Figure 4a)**, was detected in meiosis II **(Figure S3d)**. Hence we could not assess pRec8 staining in the centromere region in meiosis I when sister kinetochores are fused[20] with this antibody because we could not distinguish the specific from the unspecific signal at that location. Together with the fact that 80 % of centromeric Rec8 are removed in meiosis I already (**Figure 4a**) we thus decided to limit our analysis to arm cohesin, as the pattern of cross-shaped staining is only observed for proteins of the cohesin complex. It is however interesting to note that Rec8 seems to be phosphorylated in meiosis II as well **(Figure S3d)**, suggesting that pS482 and/or pS479 are involved in centromeric cohesin removal at anaphase II as well.

If endogenous Rec8 were phosphorylated in an Aurora B/C-dependent manner on S482 (recognized by our phospho-specific antibody), detection of phosphorylated Rec8 on arms should fail in oocytes treated with Aurora B/C inhibitor. Indeed, treatment with Aurora B/C inhibitor (ZM447439) from GVBD (germinal vesicle breakdown) onwards led to complete loss of chromosome arms staining clearly positive for Rec8 pS482 on control spreads **(Figure 4e)**. Together with the failure to correctly segregate chromosomes in meiosis I in absence of Aurora B/C activity we conclude that mouse Rec8 is phosphorylated in an Aurora B/C kinase-dependent manner on S482 for Separase cleavage in meiosis I.

The Aurora B/C consensus motif we identified here in mouse oocytes can also be found in human Rec8, suggesting conservation of the mechanism for Rec8 cleavage in mammalian oocytes. However, we do not exclude potential roles for other kinases, such as CK1δ or DDK, which are required for Rec8 cleavage in yeast[6-9] in this process. Also, the role of Plk1 in cohesin removal in oocytes has to be clarified. Future work will aim at identifying additional Rec8 kinases, their interdependencies, how Rec8 accessibility is regulated, and the roles these kinases play in the cleavage of distinct Rec8 fractions. Chromosome segregation in human oocytes is highly error prone, one reason being weakening of cohesin complexes with age, causing high rates of pregnancy loss[35]. Insights into the molecular mechanisms of cohesin removal will be crucial to better understand how these errors occur.

## Acknowledgements

We thank members of the MOM team for discussion and helpful suggestions, Eileen Breunig for preliminary experiments, Olaf Stemmann for Rad21 sensor construct, mRec8 39A plasmid and hosting EN in his lab for several months, Kim Nasmyth for the Separase conditional knock-out mouse strain, Scott Keeney for Rec8 polyclonal antibody and Léa Lemaire (Covalab) for useful advice on the generation of phospho-specific antibodies. EN was supported by a postdoctoral fellowship by the FRM (Fondation de la Recherche Médicale SPF20150934093) and EMBO (short term fellowship ASTF 426 – 2015). The KW lab obtained funding for this project by ANR (ANR-16-CE92-0007-01, ANR-19-CE13-0015), FRM (Equipe FRM DEQ 20160334921, Equipe FRM DEQ 202103012574), and core funding by the CNRS and Sorbonne Université.

## Author contributions

Most experiments in Figure 1, Figure 2, Figure 3 and Figure S3 were performed by EN, in Figure 3c and S2 by SAT, in Figure 1b by EB and in the remaining Figure panels by SEJ with contributions from CF and DC. DC additionally provided expert technical help throughout the project. Statistical analysis was done by EN, EB, SAT, SEJ and CF, figures were prepared by EN, SEJ, CF, EB, SAT and KW. Overall supervision, funding acquisition and project administration were done by KW. The manuscript was written by KW with substantial input from all authors.

## Material and Methods

### Animals

Adult CD-1 mice were purchased from Janvier, France. Separase^-/-^ mice and litter mate control mice were bred in the conventional animal facility of UMR7622 (authorization C75-05-13) with food and water access *ad libitum*. Mice were sacrificed by cervical dislocation between 8-16 weeks of age, ovaries were dissected and GV stage oocytes harvested. Mice were not involved in any procedures before sacrifice except for genotyping.

### Mouse oocyte and embryo culture

Oocytes were harvested in M2 medium (homemade or Merck Millipore, MR-015P) and kept arrested at GV stage with 100μg/mL dbcAMP (dibutyryl cyclic AMP, Sigma-Aldrich, D0260). After release in dbc-AMP free M2 medium, oocytes undergoing GVBD no later than 90min after release were used for experiments. Oocytes were cultured in M2 medium drops covered by mineral oil, manipulated on heating plates and kept in a temperature-controlled incubator at 38°C. For longer incubation, oocytes were cultured in M16 medium (MERK Millipore MR-10P or homemade) in a CO_2_- and temperature-controlled incubator. To obtain 8-cell stage embryos in mitosis, superovulation was induced in female mice by intraperitoneal injection of PMSG (pregnant mare serum gonadotropin, 5 IU/female) and hCG (human chorionic gonadotropin, 5 IU/female) 48 hours later. Then, females were coupled with a male, verified by the presence of a vaginal plug. Pregnant female mice were sacrificed 12h after fertilization, embryos collected and incubated in EmbryoMax CZB medium (Sigma-Aldrich) with 1mM supplementary L-glutamine (Sigma-Aldrich) until they reached 8-cell stage, then incubated with 4 μM Nocodazole (Sigma Aldrich) for 6 hours to keep the cells arrested in the 4th mitosis.

### Drug treatments

All inhibitors used were dissolved in DMSO (Sigma-Aldrich, D2650) and diluted to M2 or M16 medium at the indicated final concentrations. For Aurora B/C inhibition, ZM447439 (Tocris, 2458) was used at 20 μM and AZD1152 (Sigma Aldrich, SML0268) at 10μM. For Aurora A inhibition, MLN 8237 (Bertin Bio-reagent, 13602) was used at 1μM. To bypass the SAC, Mps1 inhibitor Reversine (Cayman Chemical, 10004412) was used at 0,5 μM, and also added to the oil covering the culture droplets.

### Plasmids

Plasmids used are described in Table S1. Rad21 sensor construct was a gift from O. Stemman (University of Bayreuth). Cloning of the plasmids was carried out using restriction enzymes (Thermo Fisher), PfuI polymerase (Promega) for PCR, and T4 ligation kit (Thermo Fisher). DNA was purified with QIAprep Spin Miniprep Kit, QIAquick PCR purification kit or QIAquick Gel Extraction kit (Qiagen). Cloning strategies for each construct are described in Table S1. All primer sequences are available upon request. All primers were synthesized commercially (Eurofins). After cloning, plasmid sequences were verified through a commercial sequencing service (GATC).

### Microinjection

mRNA coding for Securin-YFP, or sensor constructs was transcribed using the mMessage Machine T3 or T7 Kit (Invitrogen) and purified with RNeasy mini-kit (Qiagen). GV oocytes were injected on an inverted Nikon Eclipse Ti microscope with self-made micropipettes (Narishige magnetic puller, PN-31) and a FemtoJet microinjector pump (Eppendorf) with constant flow. Oocytes were manipulated using a holding pipette (vacuTip from Eppendorf).

### Chromosome spreads

After removal of the zona pellucida by successive transfer in acidic tyrode solution, chromosomes were spread by transferring oocytes in hypotonic solution (3 mM DTT, 0,15% Triton-X100, 1% Paraformaldehyde), placed on 8 wells glass slides(Fisher Scientific, 10588681)[36]. Upon drying at room temperature, chromosome spreads were stored at -20°C until immunostaining. Thawed spreads were washed several times with PBS 1X, incubated 10 min with PBS-BSA 3%, and incubated with primary antibodies diluted in PBS-BSA 3% for 2-3h. The following antibodies were used at the indicated concentrations: Human CREST serum (Immunovision, HCT-100, at 1:50 or 1:100), Rabbit poly clonal anti-Rec8 (gift from Scott Keeney, 1:50), Rabbit anti-pRec8 (serum A, B, C, D, E, F, 1:20), anti-phospho-Histone H3 (pS10H3, 1/100 dilution; Merck Millipore, 05-806). Spreads were washed several times with PBS and incubated with the following secondary antibodies diluted in PBS-BSA 3%, for at least 1 hour: donkey anti-human Cy3 (709-166-149, Jackson Immuno Research, 1:200), donkey anti-Human Alexa Fluor 488 (709-546-149, Jackson Immuno Research, 1:200), donkey anti-Rabbit Cy3 (711-166-152, Jackson Immuno Research, 1:200), donkey anti-Rabbit Alexa Fluor 488 (711-546-152, Jackson Immuno Research, 1:200). DNA was stained with Hoechst 33342 (Invitrogen, H21492, 50μg/mL). After several PBS washing steps chromosome spreads were mounted in Vectashield (Eurobio H-1000) or AF1 mounting medium (CityFluor) and stored at 4°C until acquisitions with a spinning disk confocal microscope.

### Image Acquisitions

Live imaging of oocytes expressing Securin-YFP was performed on a Nikon eclipseTE 2000-E inverted microscope coupled to a Prime sCMOS camera, equipped with PrecisExite High Power LED Fluorescence, a Märzhäuser Scanning Stage and using a Plan APO (20x/0,75NA) objective. Time lapse was set up for 6h with 20 min intervals. 1 z-section was acquired for every time point in the GFP channel and in DIC. Live imaging of oocytes expressing sensor constructs was performed on a Zeiss Axiovert 200M inverted spinning disk confocal microscope, coupled to an EMCD camera (Evolve 512, Photometrics), using a Plan-APO (63x/1,4 NA) oil objective. Time lapse was set up for 10 hours with 20 min intervals. For each timepoint, 11 z-stacks (3 μm optical section spacing) were acquired in 491nm channel for GFP or YFP and in 561nm channel for mCherry. All live imaging was done under perfectly temperature-controlled conditions, and microscopes were piloted by Metamorph software. Fixed chromosome spreads were acquired on the same Zeiss Axiovert 200M inverted spinning disk confocal microscope, but using a 100x/1.4 NA) oil objective. 6 z sections of 0,4 μm interval were taken in each channel. Images were processed using Fiji software. For publication, images were treated in Adobe Photoshop to adjust brightness and contrast, all adjustments were applied to whole images uniformly.

### Quantification and statistical analysis

Quantifications were performed with Fiji software. For quantification on chromosome spreads, the mean fluorescence intensities were measured on sum-projected images. A square is drawn around centromeres as depicted on the scheme in Figure 4A or on chromosome arm. The measured signal is corrected to background. Data plots and statistical analysis were obtained with PRISM6 software.

Quantifications of cleavage efficiencies of different sensors were done as described[26]. In brief, mean fluorescence signal for YFP (or GFP) and mCherry was measured in the box surrounding chromosomes during several timepoints before, during and after anaphase I, corrected for background fluorescence. The ratio between YFP (or GFP) and mCherry signals was calculated and the difference between pre-anaphase value (normalized to 100%) and first post-anaphase value derived.

---

**Supplementary Figure 1.**
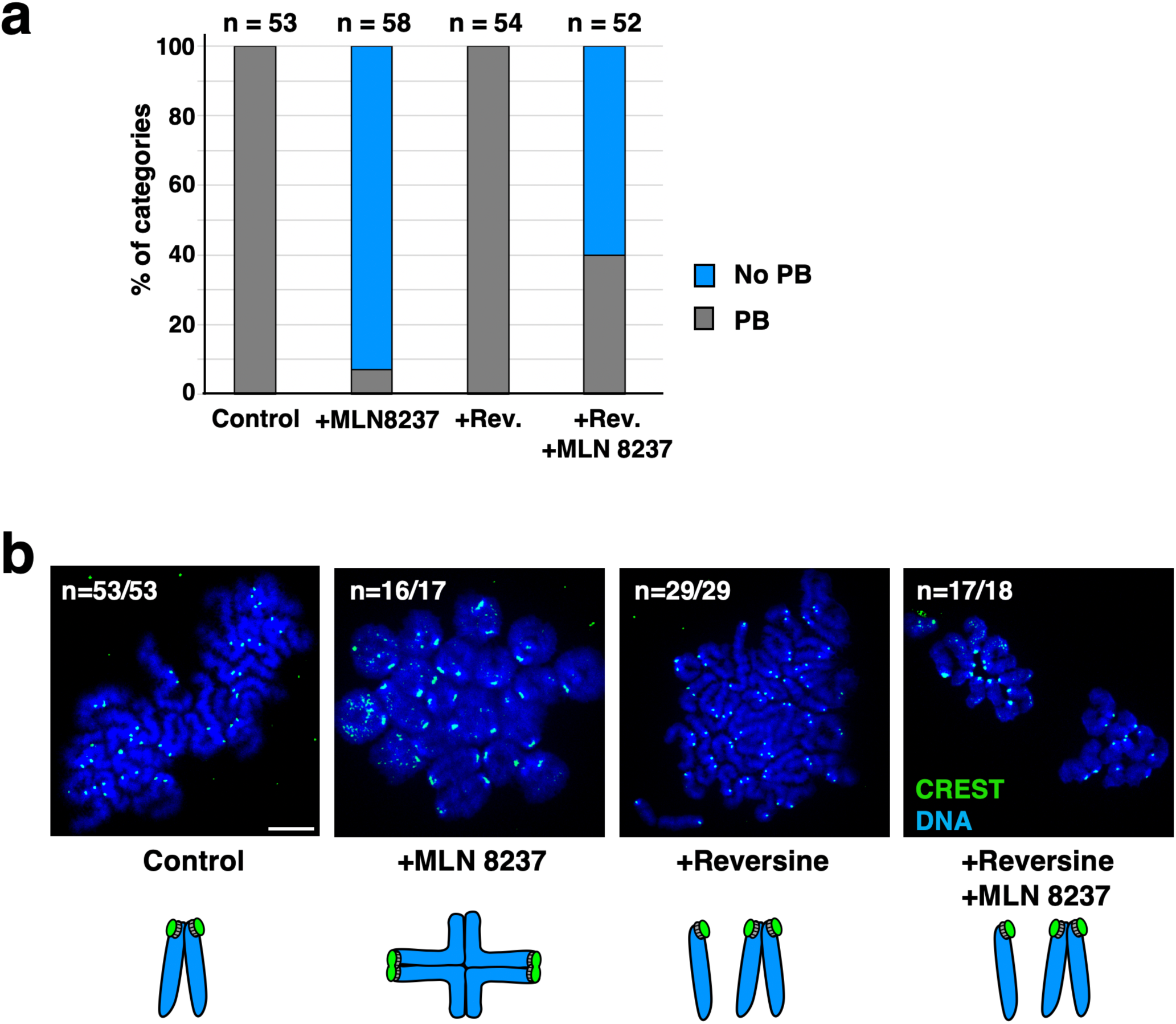
Aurora A is not required for cohesin removal in meiosis I. **a)** Oocytes were treated with 1*μ*M Aurora A inhibitor MLN8237 from GVBD onwards, and with 0,5 nM Reversine (Rev., inhibitor of the SAC kinase Mps1) in metaphase I, such as indicated. Percentage of polar body (PB) extrusion is indicated for the number of oocytes (n) observed. **b)** Chromosome spreads of oocytes at GVBD+16 hours (metaphase II in control oocytes), stained with CREST serum (centromeres, green) and Hoechst (DNA, in blue). Only oocytes that extruded PBs were analysed for MLN8237 treated oocytes. Chromosome figures are schematized below (dyads in control metaphase II oocytes, bivalents in MLN8237 treated oocytes, dyads and single sister chromatids in Reversine as well as Reversine and MLN8237 treated oocytes that extruded PBs). n indicates the number of oocytes analysed, scale bars are 10 *μ*M. Related to Figure 1

**Supplementary Figure 2.**
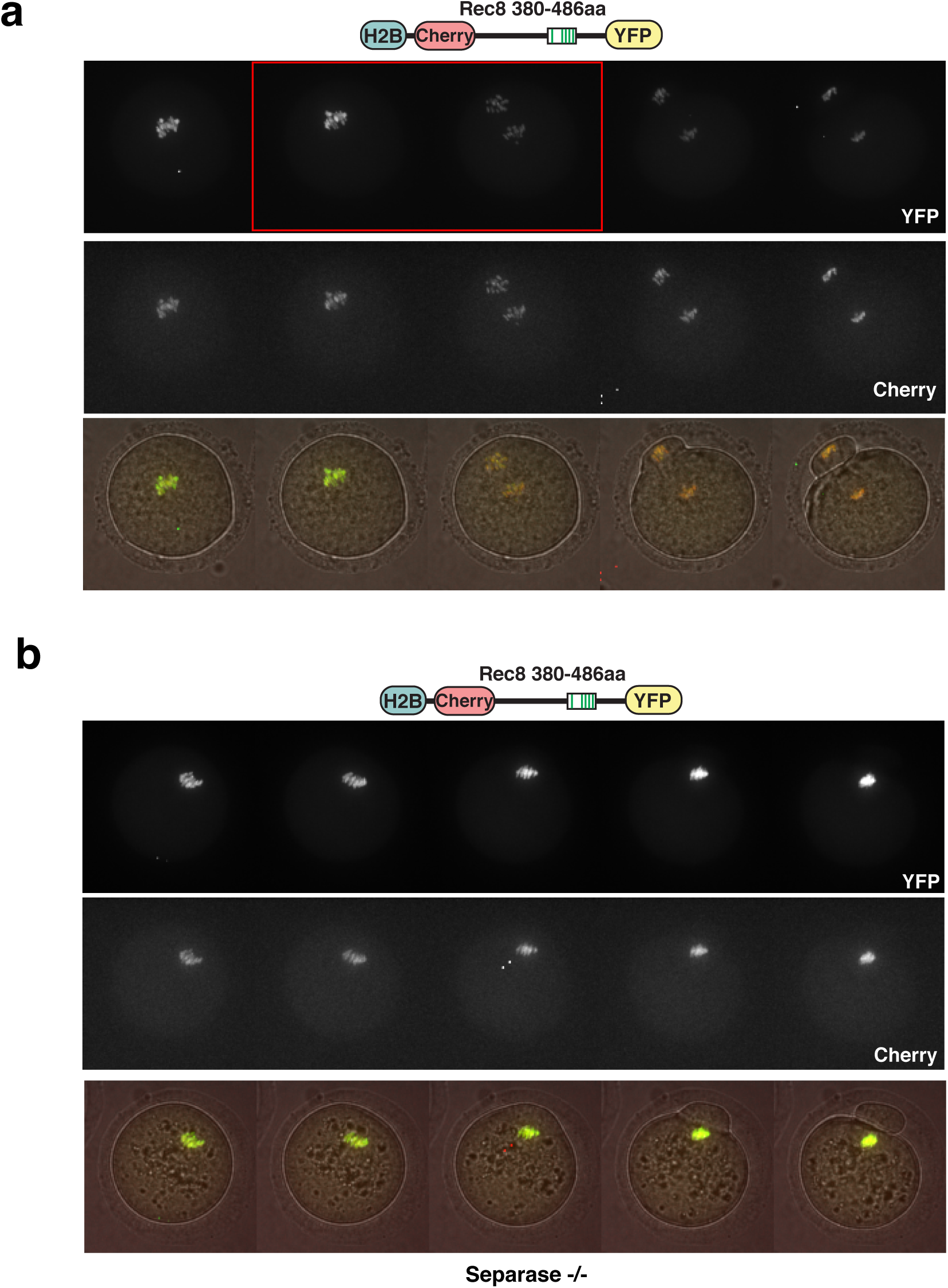
Rec8 360-486aa sensor cleavage in oocytes depends on Separase. **a, b)** Stills of time lapse movies analysed in Figure 3c), to determine cleavage efficiencies of Rec8 380-486aa sensor construct in a) Separase control (Sep^+/+^) and b) Separase knock-out (Sep^-/-^) oocytes. Overlays of z-sections of the red and green fluorescence channels are above, the overlays with DIC images below. Acquisitions +/- 40 minutes relative to onset of anaphase I are shown. Metaphase to anaphase transition is marked with a red rectangle in the control. Related to Figure 3

**Supplementary Figure 3.**
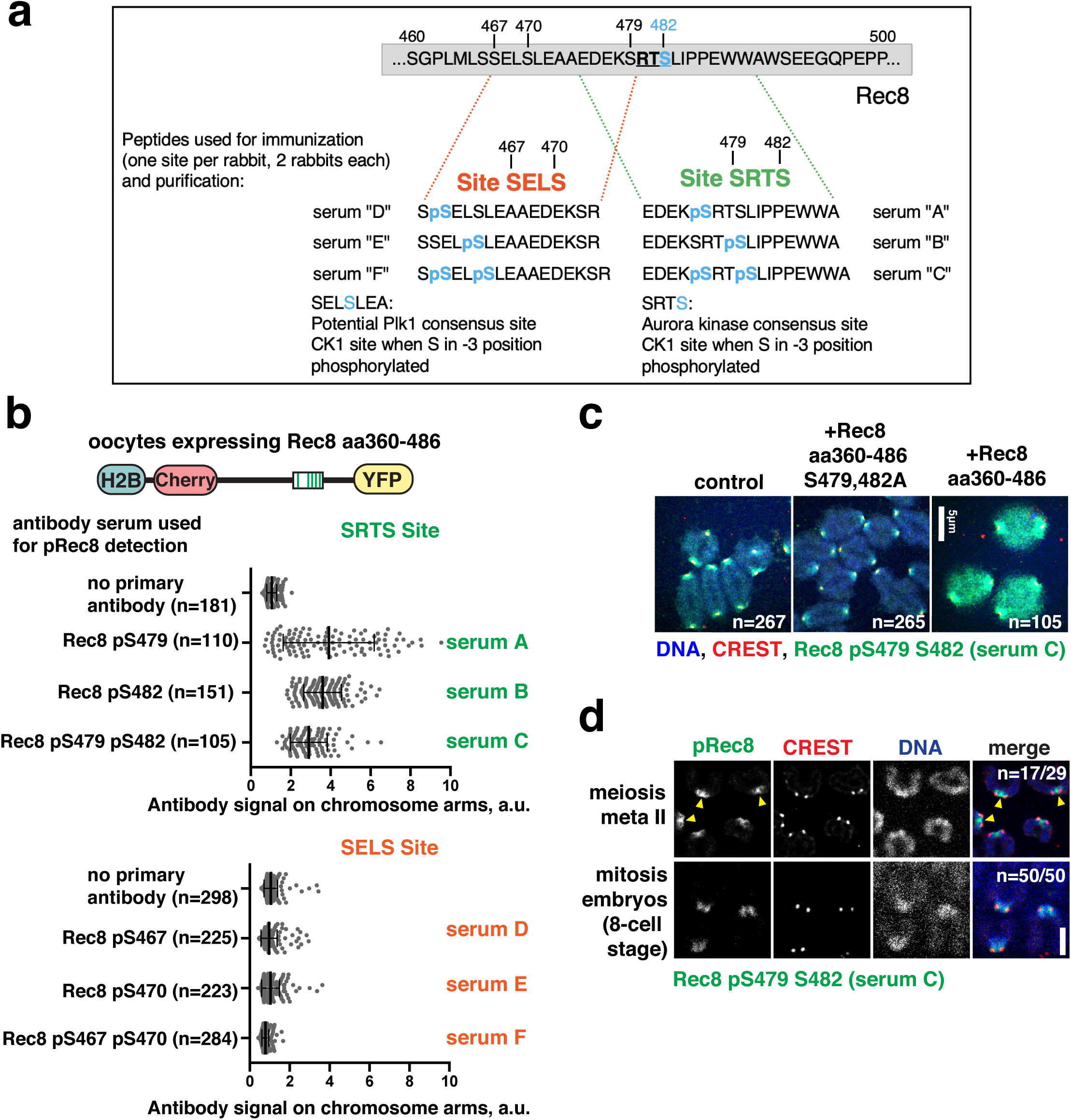
Phosphospecific antibodies recognise phosphorylated Rec8. **a)** Scheme of phospho-specific antibody generation. Two rabbits each were immunized against Rec8 peptides spanning either 466-480aa, or 475-489aa of mouse Rec8, with none, one or both Serines in positions 467, 470 or 479, 482, respectively, being phosphorylated. Sera purified against peptides carrying the indicated phosphorylated residues were designated as A-F, such as specified. n indicates number of chromosomes. **b)** Chromosome spreads of oocytes in prometaphase I (GVBD +4h) expressing the sensor construct indicated on the top were stained with the antibody sera indicated on the right. The antibody signal on chromosome arms was quantified. Each dot on the dot plots is a measurement of immunofluorescence staining of corresponding antibodies from a sister chromatid pair corrected for background. All measurements were normalized to the mean of “no primary antibody” sample. Bar and whiskers are mean and SD, a.u. – arbitrary units. n indicates number of chromosomes. **c)** Chromosome spreads of oocytes in prometaphase I (GVBD+4h) expressing the sensor construct indicated above, were stained with serum C (shown in green), centromeres with CREST (red), and DNA with Hoechst (blue). n indicates number of chromosomes. Scale bar (white) is 5*μ*m. **d)** Chromosome spreads from oocytes in metaphase II (top), or embryos at the 8-cell stage in mitosis (bottom) were stained with pRec8 serum C (green), CREST serum (centromeres, red) and Hoechst (chromosomes, blue). Yellow arrowheads indicate the signal between the two centromere dots of sister chromatids in meioisis II and not in mitosis. n indicates the number of spreads. Scale bar (white) is 5*μ*m. n indicates number of oocytes (above) ormitotic cells (below), and proportion with similar staining. Related to Figure 4

**Table S1.**

| **Construct name used in the article** | **Description** | **Mutations** | **Plasmid name** | **Cloning** |
| --- | --- | --- | --- | --- |
| negative ctrl | Sensor construct without a cleavable insert between Cherry and YFP | none | pRN3-H2B-mCherry-YFP | pRN3-H2B-mCherry-mRec8-YFP was digested with XhoI and AgeI to remove mRec8, ends were blunted with PfuI and backbone ligated. The sequence was mutated via PCR mutagenesis to correct the frame shift mutation with oligos oNE104, oNE105. |
| Rec8 wt | Sensor construct with full length wild type mouse Rec8 | none | pRN3-H2B-mCherry-mRec8-YFP | mouse Rec8 was amplified from pCS2-Flag3-Tev2-mRec8 plasmid (a gift from Olaf Stemmann) with oligos oNE44, oNE45, then subcloned into pRN3-YFP via XhoI and AgeI sites. H2B-mCherry was amplified with oligos oNE46, oNE47 and subcloned into the intermediate pRN3-Rec8-YFP plasmid via XhoI. |
| Rec8 non-ph (Rec8 39S/T-A) | Sensor construct with full length mouse Rec8 rendered non-phosphorylatable by mutating 39S/T into A | S149A T164A<br>T168A T172A<br>T177A T186A<br>T189A S192A<br>T197A S219A<br>S251A S257A<br>S271A S274A<br>T313A S325A<br>T326A S332A<br>T353A S356A<br>T366A T384A<br>S385A T392A<br>T394A S396A<br>T408A T426A<br>T429A T444A<br>S448A S460A<br>S466A S467A<br>S470A S479A<br>T481A S482A<br>S492A | pRN3-H2B-mCherry-39ARec8-YFP | Rec8 39S/T-A was subcloned from pCS2-H2B-Cherry-39AmRec8-YFP (a gift from Olaf Stemmann, 39AmRec8 was commercially synthesized) to replace wt mRec8 in pRN3-H2B-mCherry-mRec8-YFP plasmid. |
| Rec8 1-143aa | Sensor construct with mouse Rec8 1-143aa fragment | none | pRN3-H2B-mCherry-Rec8Nterm-YFP | pRN3-H2B-mCherry-mRec8-YFP was digested with BamHI and AgeI (Thermo Fisher) to remove most of Rec8 sequence except for 1-143aa. The resulting plasmid was corrected for cloning artifacts with PCR mutagenesis using oligos oNE110, oNE111. |
| Rec8 143-290aa | Sensor construct with mouse Rec8 143-290aa fragment | none | pRN3-H2B-mCherry-Rec8(143-290)-YFP | Mouse Rec8 143-290aa fragment was amplified from pRN3-H2B-mCherry-mRec8-YFP plasmid with oligos oNE100, oNE101 and subcloned to replace full-length Rec8 via XhoI and AgeI sites. |
| Rec8 380-486aa | Sensor construct with mouse Rec8 380-486aa fragment | none | pRN3-H2B-mCherry-Rec8(380-486)-YFP | Mouse Rec8 380-486aa fragment was amplified from pRN3-H2B-mCherry-mRec8-YFP plasmid with oligos oNE102, oNE103 and subcloned to replace full-length Rec8 via XhoI and AgeI sites. |
| Rec8 380-486aa ExxD*2 (Rec8 380-486aa R434D, R454D) | Sensor construct with mouse Rec8 380-486aa fragment where two potential Separase cleavage sites were disabled | R434D R454D | pRN3-H2B-mCherry-Rec8(380-486)R434D R454D -YFP | Plasmid pRN3-H2B-mCherry-Rec8(380-486)-YFP was modified through PCR mutagenesis with oligos oNE114 and oNE115, then oNE116 and oNE117. |
| Rec8 380-486aa non-ph (Rec8 380-486aa 17 S/T-A) | Sensor construct with mouse Rec8 380-486aa fragment rendered non-phosphorylatable by mutating 17S/T into A | T384A S385A T392A T394A S396A T408A T426A T429A T444A S448A S460A S466A S467A S470A S479A T481A S482A | pRN3-H2B-mCherry-Rec8(380-486)17A-YFP | Mouse Rec8(380-486aa)17A fragment was amplified from pRN3-H2B-mCherry-39AmRec8-YFP plasmid with oligos oNE112, oNE113 and subcloned to replace full-length Rec8 via XhoI and AgeI sites. |
| Rec8 380-486aa AAELA ARTA 5S-A | Sensor construct with mouse Rec8 380-486aa fragment where the potential important phosphosites were rendered non-phosphorylatable | S482A S479A S466A S467A S470A | pRN3-H2B-mCherry-Rec8(380-486) S482A S479A S466A S467A S470A-YFP | pRN3-H2B-mCherry-Rec8(380-486)-YFP was modified through several rounds of PCR mutagenesis, first with oligos oNE62 and oNE63, then oNE130 and oNE131, then oNE138 and oNE139. |

## References

1. Sato, M., Kakui, Y., and Toya, M. (2021). Tell the Difference Between Mitosis and Meiosis: Interplay Between Chromosomes, Cytoskeleton, and Cell Cycle Regulation. Front Cell Dev Biol 9, 660322.

2. Miller, M.P., Amon, A., and Unal, E. (2013). Meiosis I: when chromosomes undergo extreme makeover. Curr Opin Cell Biol 25, 687–696.

3. Keating, L., Touati, S.A., and Wassmann, K. (2020). A PP2A-B56-Centered View on Metaphase-to-Anaphase Transition in Mouse Oocyte Meiosis I. Cells 9.

4. Kitajima, T.S., Sakuno, T., Ishiguro, K., Iemura, S., Natsume, T., Kawashima, S.A., and Watanabe, Y. (2006). Shugoshin collaborates with protein phosphatase 2A to protect cohesin. Nature 441, 46–52.

5. Riedel, C.G., Katis, V.L., Katou, Y., Mori, S., Itoh, T., Helmhart, W., Galova, M., Petronczki, M., Gregan, J., Cetin, B., et al. (2006). Protein phosphatase 2A protects centromeric sister chromatid cohesion during meiosis I. Nature 441, 53–61.

6. Katis, V.L., Lipp, J.J., Imre, R., Bogdanova, A., Okaz, E., Habermann, B., Mechtler, K., Nasmyth, K., and Zachariae, W. (2010). Rec8 phosphorylation by casein kinase 1 and Cdc7-Dbf4 kinase regulates cohesin cleavage by separase during meiosis. Developmental cell 18, 397–409.

7. Rumpf, C., Cipak, L., Dudas, A., Benko, Z., Pozgajova, M., Riedel, C.G., Ammerer, G., Mechtler, K., and Gregan, J. (2010). Casein kinase 1 is required for efficient removal of Rec8 during meiosis I. Cell Cycle 9, 2657–2662.

8. Ishiguro, T., Tanaka, K., Sakuno, T., and Watanabe, Y. (2010). Shugoshin-PP2A counteracts casein-kinase-1-dependent cleavage of Rec8 by separase. Nature cell biology 12, 500–506.

9. Le, A.H., Mastro, T.L., and Forsburg, S.L. (2013). The C-terminus of S. pombe DDK subunit Dfp1 is required for meiosis-specific transcription and cohesin cleavage. Biol Open 2, 728–738.

10. Ferrandiz, N., Barroso, C., Telecan, O., Shao, N., Kim, H.M., Testori, S., Faull, P., Cutillas, P., Snijders, A.P., Colaiacovo, M.P., et al. (2018). Spatiotemporal regulation of Aurora B recruitment ensures release of cohesion during C. elegans oocyte meiosis. Nature communications 9, 834.

11. Nguyen, A.L., and Schindler, K. (2017). Specialize and Divide (Twice): Functions of Three Aurora Kinase Homologs in Mammalian Oocyte Meiotic Maturation. Trends in genetics : TIG 33, 349–363.

12. Marston, A.L., and Wassmann, K. (2017). Multiple Duties for Spindle Assembly Checkpoint Kinases in Meiosis. Front Cell Dev Biol 5, 109.

13. Ditchfield, C., Johnson, V.L., Tighe, A., Ellston, R., Haworth, C., Johnson, T., Mortlock, A., Keen, N., and Taylor, S.S. (2003). Aurora B couples chromosome alignment with anaphase by targeting BubR1, Mad2, and Cenp-E to kinetochores. The Journal of cell biology 161, 267–280.

14. Mortlock, A.A., Foote, K.M., Heron, N.M., Jung, F.H., Pasquet, G., Lohmann, J.J., Warin, N., Renaud, F., De Savi, C., Roberts, N.J., et al. (2007). Discovery, synthesis, and in vivo activity of a new class of pyrazoloquinazolines as selective inhibitors of aurora B kinase. J Med Chem 50, 2213–2224.

15. Balboula, A.Z., and Schindler, K. (2014). Selective disruption of aurora C kinase reveals distinct functions from aurora B kinase during meiosis in mouse oocytes. PLoS genetics 10, e1004194.

16. Swain, J.E., Ding, J., Wu, J., and Smith, G.D. (2008). Regulation of spindle and chromatin dynamics during early and late stages of oocyte maturation by aurora kinases. Mol Hum Reprod 14, 291–299.

17. Sharif, B., Na, J., Lykke-Hartmann, K., McLaughlin, S.H., Laue, E., Glover, D.M., and Zernicka-Goetz, M. (2010). The chromosome passenger complex is required for fidelity of chromosome transmission and cytokinesis in meiosis of mouse oocytes. Journal of cell science 123, 4292–4300.

18. Hirota, T., Lipp, J.J., Toh, B.H., and Peters, J.M. (2005). Histone H3 serine 10 phosphorylation by Aurora B causes HP1 dissociation from heterochromatin. Nature 438, 1176–1180.

19. Vallot, A., Leontiou, I., Cladiere, D., El Yakoubi, W., Bolte, S., Buffin, E., and Wassmann, K. (2018). Tension-Induced Error Correction and Not Kinetochore Attachment Status Activates the SAC in an Aurora-B/C-Dependent Manner in Oocytes. Curr Biol 28, 130–139 e133.

20. Gryaznova, Y., Keating, L., Touati, S.A., Cladiere, D., El Yakoubi, W., Buffin, E., and Wassmann, K. (2021). Kinetochore individualization in meiosis I is required for centromeric cohesin removal in meiosis II. The EMBO journal, e106797.

21. Nguyen, A.L., Drutovic, D., Vazquez, B.N., El Yakoubi, W., Gentilello, A.S., Malumbres, M., Solc, P., and Schindler, K. (2018). Genetic Interactions between the Aurora Kinases Reveal New Requirements for AURKB and AURKC during Oocyte Meiosis. Curr Biol 28, 3458–3468 e3455.

22. Saskova, A., Solc, P., Baran, V., Kubelka, M., Schultz, R.M., and Motlik, J. (2008). Aurora kinase A controls meiosis I progression in mouse oocytes. Cell Cycle 7, 2368–2376.

23. Blengini, C.S., Ibrahimian, P., Vaskovicova, M., Drutovic, D., Solc, P., and Schindler, K. (2021). Aurora kinase A is essential for meiosis in mouse oocytes. PLoS genetics 17, e1009327.

24. Manfredi, M.G., Ecsedy, J.A., Chakravarty, A., Silverman, L., Zhang, M., Hoar, K.M., Stroud, S.G., Chen, W., Shinde, V., Huck, J.J., et al. (2011). Characterization of Alisertib (MLN8237), an investigational small-molecule inhibitor of aurora A kinase using novel in vivo pharmacodynamic assays. Clin Cancer Res 17, 7614–7624.

25. Touati, S.A., Buffin, E., Cladiere, D., Hached, K., Rachez, C., van Deursen, J.M., and Wassmann, K. (2015). Mouse oocytes depend on BubR1 for proper chromosome segregation but not for prophase I arrest. Nature communications 6, 6946.

26. Nikalayevich, E., Bouftas, N., and Wassmann, K. (2018). Detection of Separase Activity Using a Cleavage Sensor in Live Mouse Oocytes. Methods Mol Biol 1818, 99–112.

27. Shindo, N., Kumada, K., and Hirota, T. (2012). Separase sensor reveals dual roles for separase coordinating cohesin cleavage and cdk1 inhibition. Developmental cell 23, 112–123.

28. Burkhardt, S., Borsos, M., Szydlowska, A., Godwin, J., Williams, S.A., Cohen, P.E., Hirota, T., Saitou, M., and Tachibana-Konwalski, K. (2016). Chromosome Cohesion Established by Rec8-Cohesin in Fetal Oocytes Is Maintained without Detectable Turnover in Oocytes Arrested for Months in Mice. Curr Biol 26, 678–685.

29. Maier, N.K., Ma, J., Lampson, M.A., and Cheeseman, I.M. (2021). Separase cleaves the kinetochore protein Meikin at the meiosis I/II transition. Developmental cell 56, 2192–2206 e2198.

30. Kudo, N.R., Anger, M., Peters, A.H., Stemmann, O., Theussl, H.C., Helmhart, W., Kudo, H., Heyting, C., and Nasmyth, K. (2009). Role of cleavage by separase of the Rec8 kleisin subunit of cohesin during mammalian meiosis I. Journal of cell science 122, 2686–2698.

31. Kudo, N.R., Wassmann, K., Anger, M., Schuh, M., Wirth, K.G., Xu, H., Helmhart, W., Kudo, H., McKay, M., Maro, B., et al. (2006). Resolution of chiasmata in oocytes requires separase-mediated proteolysis. Cell 126, 135–146.

32. Ogushi, S., Rattani, A., Godwin, J., Metson, J., Schermelleh, L., and Nasmyth, K. (2020). Loss of Sister Kinetochore Co-orientation and Peri-centromeric Cohesin Protection after Meiosis I Depends on Cleavage of Centromeric REC8. bioRxiv.

33. Kumar, M., Gouw, M., Michael, S., Samano-Sanchez, H., Pancsa, R., Glavina, J., Diakogianni, A., Valverde, J.A., Bukirova, D., Calyseva, J., et al. (2020). ELM-the eukaryotic linear motif resource in 2020. Nucleic Acids Res 48, D296–D306.

34. Tachibana-Konwalski, K., Godwin, J., van der Weyden, L., Champion, L., Kudo, N.R., Adams, D.J., and Nasmyth, K. (2010). Rec8-containing cohesin maintains bivalents without turnover during the growing phase of mouse oocytes. Genes & development 24, 2505–2516.

35. Herbert, M., Kalleas, D., Cooney, D., Lamb, M., and Lister, L. (2015). Meiosis and Maternal Aging: Insights from Aneuploid Oocytes and Trisomy Births. Cold Spring Harb Perspect Biol 7.

36. Chambon, J.P., Hached, K., and Wassmann, K. (2013). Chromosome spreads with centromere staining in mouse oocytes. Methods Mol Biol 957, 203–212.

